# Ectopic cambia in Japanese wisteria (*Wisteria floribunda*) vines are associated with the expression of conserved *KNOX* genes

**DOI:** 10.1101/2024.08.07.606835

**Authors:** Israel L. Cunha Neto, Anthony A. Snead, Jacob B. Landis, Chelsea D. Specht, Joyce G. Onyenedum

## Abstract

Secondary growth is a conserved mechanism that gives rise to vascular tissues produced via a single vascular cambium. Molecular mechanisms underlying this process are characterized mostly in model species bearing typical vascular architecture, while the genetics underlying ecologically-important atypical vascular architectures remain unexplored. We use developmental anatomy, comparative transcriptomics, and molecular evolutionary analyses to address this knowledge gap, investigating how multiple ectopic cambia (EC) form in the woody vine Japanese wisteria. Anatomical studies show EC in Japanese wisteria arise from cortical parenchyma, while cambium-specific transcriptome comparisons reveal that genes acting as regulators of typical cambium development in model species are likewise associated with atypical EC development. Gene trees of KNOX proteins indicate duplication events may contribute to EC formation, including a Fabaceae-specific duplication of KNAT6 detected as under positive selection. These findings reveal insights into the genetics of EC formation, advancing our understanding of the development of complex vascular traits.

## Introduction

The vast majority of woody plants undergo a common type of secondary growth that involves the production of a single meristematic niche called the vascular cambium, responsible for generating secondary xylem (wood) and secondary phloem (inner bark) and enabling the plants to expand in diameter^1^. With only a single vascular cambium, plants can live up to 5,000 years (e.g. Bristlecone pines [*Pinus longaeva* D.K.Bailey] or build a sophisticated plumbing system to transport water upwards of 380 ft (e.g. Coastal Redwood [*Sequoia sempervirens* (D.Don) Endl.]^2,3^. However, some plants deviate from the canonical modality to create woody structures with unique growth patterns that result in novel anatomical formations, such as those found within the stems of woody vines or “lianas”. Lianas reach the forest canopy by climbing over other plants rather than supporting their weight^4^. As they weave their way through dense forest understory canopy, the stems of lianas are frequently damaged either through the collapse of host trees, or the regular wear and tear of twisting and twining^5^. The woody stems of many lianas present with “ectopic cambia”, a developmental aberration that may be a mechanism for lianas to repair their stems after injury while maintaining key vascular functions of transportation and support^5^.

Ectopic cambia (EC) result from multiple vascular cambia developing in atypical locations, secondary to the first-formed and central vascular cambium^6^. In addition to their role in lianas, EC may also facilitate the success of self-supporting herbs, shrubs, and trees living in harsh conditions such as mangroves and arid environments^7,8^. EC have been identified in the stems of the temperate *Wisteria* vines^9,10^, potentially facilitating their success as invasive vines in the Eastern United States^11^.

To date, all anatomical studies show that EC arise through three steps: (1) de-differentiation of living parenchyma cells, (2) cell division to give rise to a new cambium, and (3) production and differentiation of localized vascular tissues from the new cambium layer^7,9,12^. While the vast majority of the literature covering the natural diversity of EC is purely anatomical and descriptive, transgenic phenotypes in model species suggest that ECs can emerge by simply manipulating the expression of conserved genes responsible for vascular development^6^.

Recent investigations demonstrate that *Arabidopsis thaliana* (L.) Heynh and *Populus* spp. share a suite of genes in the regulatory pathway controlling the canonical model of secondary growth by a single vascular cambium^1,13,14^. Notably, transgenic lines that perturb the expression of conserved vascular genes produce EC in stems and/or roots^6^. Various studies indicate that EC can appear as the result of the misexpression of members of the class III homeodomain-leucine zipper (HD-ZIP III) transcription factor genes: for example, EC are developed in *Populus* stems through overexpression of *PRE*, an ortholog of the gene *REVOLUTA (REV)*^15^, while EC have been shown to develop in *A. thaliana* through the overexpression of *AtHB7*^16^ and repression of *AtHB4*^17^. ECs can also be formed following overexpression of *WOX4*, a member of the WOX gene family^18^, and EC were observed in double mutants following the overexpression of *WOX4* and repression of *PETAL LOSS* (*PTL*)^19^. Finally, regenerated cambium can be induced by girdling trees, allowing investigators to characterize how such cambia (here categorized as EC given their de novo establishment) can function to stitch together a wounded vascular cambium^20,21^.

KNOTTED1-like HOMEOBOX transcription factors––key regulators of vascular proliferation––often emerge as differentially expressed genes in cambium development studies^16,17,19,22^. Yet, their role is underexplored in the formation of EC development. Our current knowledge of EC formation arises from molecular manipulation or direct perturbation––either mechanical (i.e., girdling) or genetic (i.e., transgenics)––in model species that don’t normally form EC. To understand how EC contribute to plant diversity in form and function, we must understand how EC are produced in systems in which they naturally occur, particularly in lianas where they have a key functional role in allowing plants to climb.

In this work, we seek to elucidate the developmental anatomy, transcriptomic profile, and molecular evolution underlying EC formation in a pair of vines in the Fabaceae family, the ectopic cambia-producing Japanese wisteria (*Wisteria floribunda* (Willd.) DC.) and the non-ectopic cambia producing Common bean (*Phaseolus vulgaris* L.). We confirm previous reports that ectopic cambia form within the cortex of Japanese wisteria in a haphazard fashion. To reveal the transcriptomic profile underlying EC formation, we leveraged tangential cryosectioning to isolate tissues of interest for RNA sequencing. Here we report the differential expression of several conserved genes in species and cross-species analyses of cambium-specific samples, including known cambium markers and genes involved in epigenetic regulation. We identified *KNOX* genes as prospective candidates in EC development in Japanese wisteria, as multiple transcript clusters are differentially expressed in typical v. ectopic cambia contrasts. We adopted a phylogenetic approach to investigate the molecular evolution of *KNOX* genes across 45 species spanning gymnosperms and angiosperms. We implemented branch models to investigate the ratio of nonsynonymous to synonymous substitutions and assess the signatures of selection during the evolution and diversification of the *KNOX* protein family. Our analyses uncovered transcript clusters for *KNOX* proteins under positive selection, further corroborating their important role in EC development in Japanese wisteria. Together, these results expand our understanding of both typical and atypical cambium development, while identifying promising contenders underpinning EC formation in woody plants.

## Results

### Common bean and Japanese wisteria exhibit typical cambium development; Japanese wisteria develops ectopic cambia during the later stages of stem growth

To characterize the origin of ectopic cambia (EC), we began by detailing the developmental anatomy of the Common bean and Japanese wisteria (Fig. 1a, e). Young stems of both Common bean and Japanese wisteria have the same organization of primary tissues, with vascular bundles arranged in a ring (Fig. 1b, f). In young stems (3^rd^ to 5^th^ internodes, see Fig. 1a, e), we find the initiation of the vascular cambium from the fascicular cambium within each bundle and the interfascicular cambium connecting the bundles (Fig. 1c, g), which gives rise to secondary xylem (wood) inward and secondary phloem (inner bark) outward (Fig. 1d, h). This latter stage marks the final step in stem development of Common bean (Fig. 1d) giving rise to the mature form; however, Japanese wisteria development continues with further elaborations (Fig. 1i) to arrive at maturity. In stems with at least 20mm diameter, Japanese wisteria starts to produce additional cambia ectopically (Fig. 1j), with older stems displaying multiple layers of EC and its products, i.e., secondary xylem and secondary phloem (Fig. 1i). EC in Japanese wisteria arise from the cortex (Fig. 1j) where existing cortical cells undergo localized cell divisions, creating a mass of cells from which a new meristem will arise to produce secondary xylem and phloem (Fig. 1j). These EC do not arise in a uniform concentric pattern but rather initiate asynchronously across the stem circumference forming discrete strands of secondary xylem and secondary phloem (Fig. 1i).

**Fig 1.**
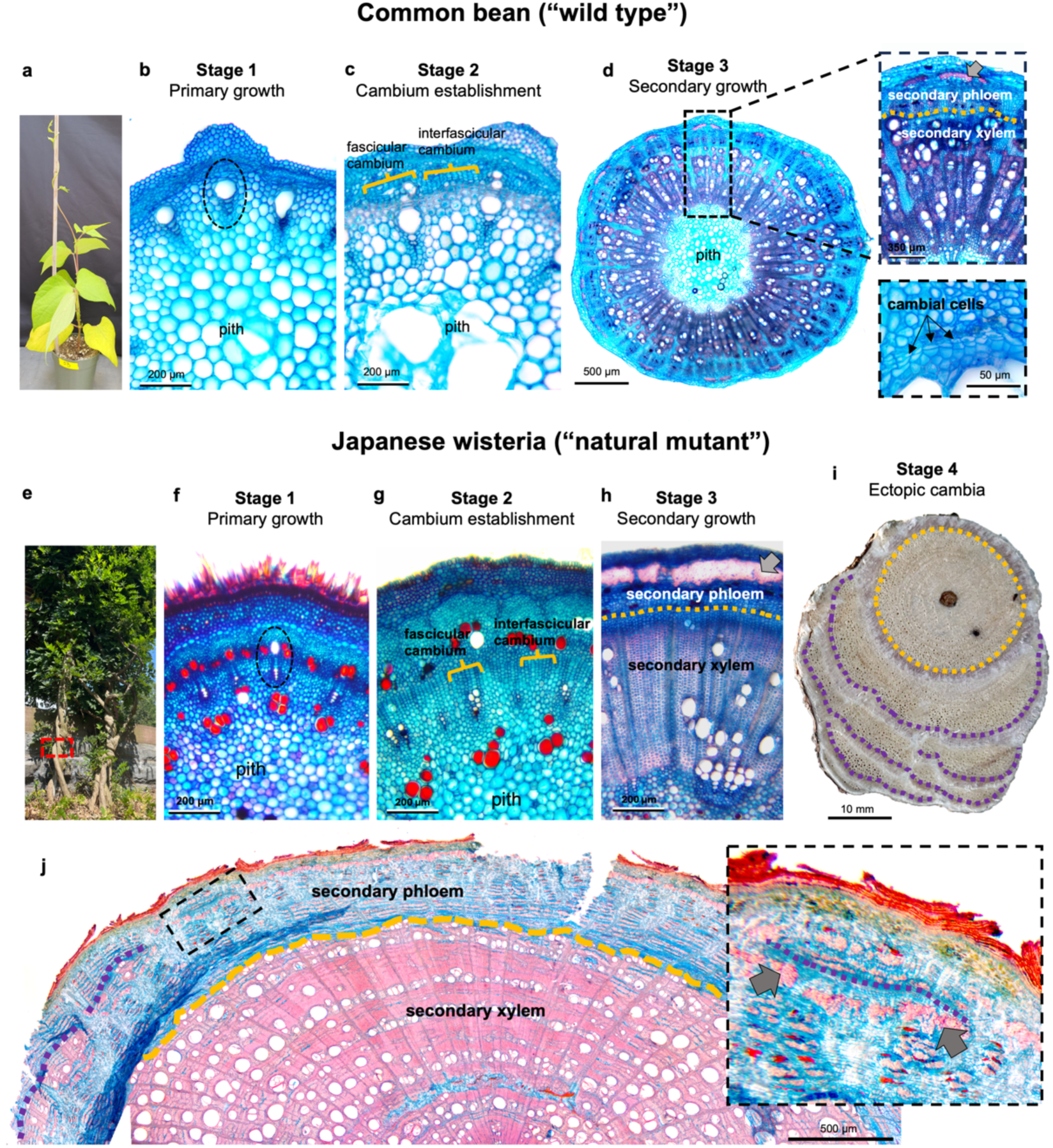
Vascular development in Common bean (*Phaseolus vulgaris*) and Japanese wisteria (*Wisteria floribunda*) share conserved developmental stages, while Japanese wisteria later forms ectopic cambia (EC) from cortical parenchyma. **a-d** Common bean. **e-i** Japanese wisteria. **a, e** Common bean and Japanese wisteria plants sampled for developmental anatomy. The dashed rectangle in “**e**” indicates the position of the image displayed in stage 4 (**i**). **b, f** Primary growth showing vascular bundles (dashed circle), formed by xylem inwards and phloem outwards, arranged in a single ring delimiting the pith. **c, g** Initiation of typical cambium from the fascicular cambium located within the bundles, and the interfascicular cambium between the bundles. **d, h** Secondary growth generated by the typical cambium (dashed yellow line) producing secondary xylem and secondary phloem. In **d,** the bottom right inset shows details of the typical cambium extracted from the top right inset. **i,** Mature stem of Japanese wisteria with EC (dashed purple line) forming multiple vascular increments in addition to the vascular cylinder produced by the typical cambium (dashed yellow line). **j** Detail of stem in advanced typical secondary growth derived from the typical cambium (dashed yellow line) and the emergence of EC (dashed purple line) in Japanese wisteria; the dashed box indicates the position of the inset (on the right). Grey arrow = pericyclic fibers. All figures are displayed in cross-section and indicate magnification.

### Transcriptomes of Common bean and Japanese wisteria

Having isolated key developmental stages, we sought to identify which genes underlie the formation of EC. Transcriptomic profiles were made from the typical cambium from the Common bean and Japanese wisteria and the EC of Japanese wisteria, with at least five biological replicates for each of the three tissues sampled. Sequencing resulted in an average of 21,420,218 (128,521,310 total) and 23,468,348 (258,151,832 total) paired-end reads per sample for the Common bean and Japanese wisteria, respectively. The transcriptome assemblies vary in BUSCO, mapping rate, and N50, but both assemblies resulted in BUSCO scores and mapping rates over 80% with N50s greater than 1500 base pairs (bp) (Table 1).

**Table 1.** The transcriptome assembly statistics (i.e., BUSCO, Mapping Rate, and N50) for the final transcriptome assemblies for *Phaseolus vulgaris* (Common bean) and *Wisteria floribunda* (Japanese wisteria).

| Species | BUSCO | Mapping Rate | N50 |
| --- | --- | --- | --- |
| Bean | C:81.8%[S:77.1%,D:4.7%], F:3.9%, M:14.3% | 82.41% | 1562 |
| Wisteria | C:88.4%[S:79%,D:9.4%], F:1.6%, M:10% | 84.99% | 1624 |

### Differential Cluster Expression and Gene Ontology Enrichment

#### Comparison across species

Using the transcript clusters for Common bean (42,827) and Japanese wisteria (74,960), 10,498 one-to-one orthologous transcript clusters were identified and passed filtering (Expressed > 0.5 counts per million). The number of differentially expressed clusters varied dramatically between contrasts. The most differentially expressed (DE) transcript clusters were found in the “Japanese wisteria v. Common bean” contrast, representing overall differences between the two species (> 6,000). This was followed by the number of DE transcripts when comparing the “Typical Cambia: Japanese wisteria v. Common bean” (> 5,000). The third highest number of DE transcripts was found in the “Ectopic Cambia v. Typical Cambia of Japanese wisteria and Common bean” contrast (> 2,000) (Table 2). Lastly, we compared the “Japanese wisteria: Ectopic v. Typical cambia” contrast, which surprisingly yielded no DE transcript clusters in this analysis (see further analyses “*Comparisons within Japanese wisteria*” below). In annotating the DE transcript clusters with Trinotate^23^ against the UniProt database for Fabales (UniProt Taxon ID 72025^24^), we found transcript clusters related to DNA methylation (e.g., methyltransferases, DNA methylase), auxin (e.g., indoleacetic acid, auxin response factors), cell cycle control (e.g., cyclins, elongation effective 1), cell differentiation (e.g., *KNOX*) and other cambial markers (e.g., *PXY*). The “Ectopic Cambia v. Typical Cambia of Japanese wisteria and Common bean” contrast is the only comparison that yielded enriched gene ontology terms based on identified DE clusters, and the GO terms are mostly related to RNA binding, splicing, and processing (Table 3).

**Table 2.** Significantly differentially expressed transcript clusters based on a false discovery rate of less than 0.05 for each contrast (one-to-one orthologous analysis).

| Contrast | Direction | Genes |
| --- | --- | --- |
| Japanese wisteria v. Common bean | Up | 3,168 |
|  | Down | 3,071 |
| Typical Cambia: Japanese wisteria v. Common bean | Up | 2,832 |
|  | Down | 2,594 |
| Ectopic Cambia v. Typical Cambia of Japanese wisteria and Common bean | Up | 1,636 |
|  | Down | 730 |

**Table 3.** The significantly enriched gene ontology (GO) terms (< 0.05 FDR) for differentially expressed transcript clusters in the “Ectopic Cambia v. Typical Cambia of Japanese wisteria and Common bean” contrast.

| Term | Gene Ontology | Direction | p-value |
| --- | --- | --- | --- |
| mRNA processing | Biological Process | ▲ | 0.0002 |
| mRNA Splicing, via Spliceosome | Biological Process | ▲ | 0.0007 |
| RNA Splicing | Biological Process | ▲ | 0.04 |
| UV Protection | Biological Process | ▲ | 0.04 |
| Spliceosomal Complex | Cellular Component | ▲ | 0.002 |
| Nucleolus | Cellular Component | ▲ | 0.004 |
| Nucleus | Cellular Component | ▲ | 0.006 |
| RNA binding | Molecular Function | ▲ | 0.000005 |
| mRNA binding | Molecular Function | ▲ | 0.003 |
| nucleic acid binding | Molecular Function | ▲ | 0.04 |
| DNA-directed 5'-3' RNA polymerase activity | Molecular Function | ▲ | 0.05 |

Footnote: Each term includes its ontology and corrected p-value rounded to one significant figure. “Ectopic Cambia v. Typical Cambia of Japanese wisteria and Common bean” is the only contrast with enriched GO terms using the differentially expressed transcript clusters. The filled arrows indicate if the terms were significantly overrepresented (▲).

##### Comparison within Japanese wisteria

Given our focus on elucidating the genes related to ectopic cambia, we narrowed our focus on genes differentially expressed in the “Japanese wisteria: Ectopic v. Typical cambia” contrast. This approach enabled us to isolate the most likely candidate genes associated with EC formation in this species. After filtering (Expressed > 0.5 counts per million in at least two samples), 46,225 transcript clusters were retained for differential transcript cluster expression analysis across the two Japanese wisteria cambium types using dream^25^, with 14 clusters (2▴ and 12▾) found to be differentially expressed. The differentially expressed clusters include transcripts related to DNA methylation, cell expansion (e.g., expansin-like B1), and cell differentiation (e.g., *KNOX*), all of which were found to be downregulated in the EC of Japanese wisteria (Fig. 2; Supplementary Table 1). Only two transcript clusters are upregulated, one of which includes the epigenetic gene Methyltransf_7 (Table 4). Interestingly, we found five differentially expressed *KNOX* genes across the four contrasts (Table 5), which makes them of particular interest given their defined role in vascular cambium proliferation in model systems.

**Fig. 2.**
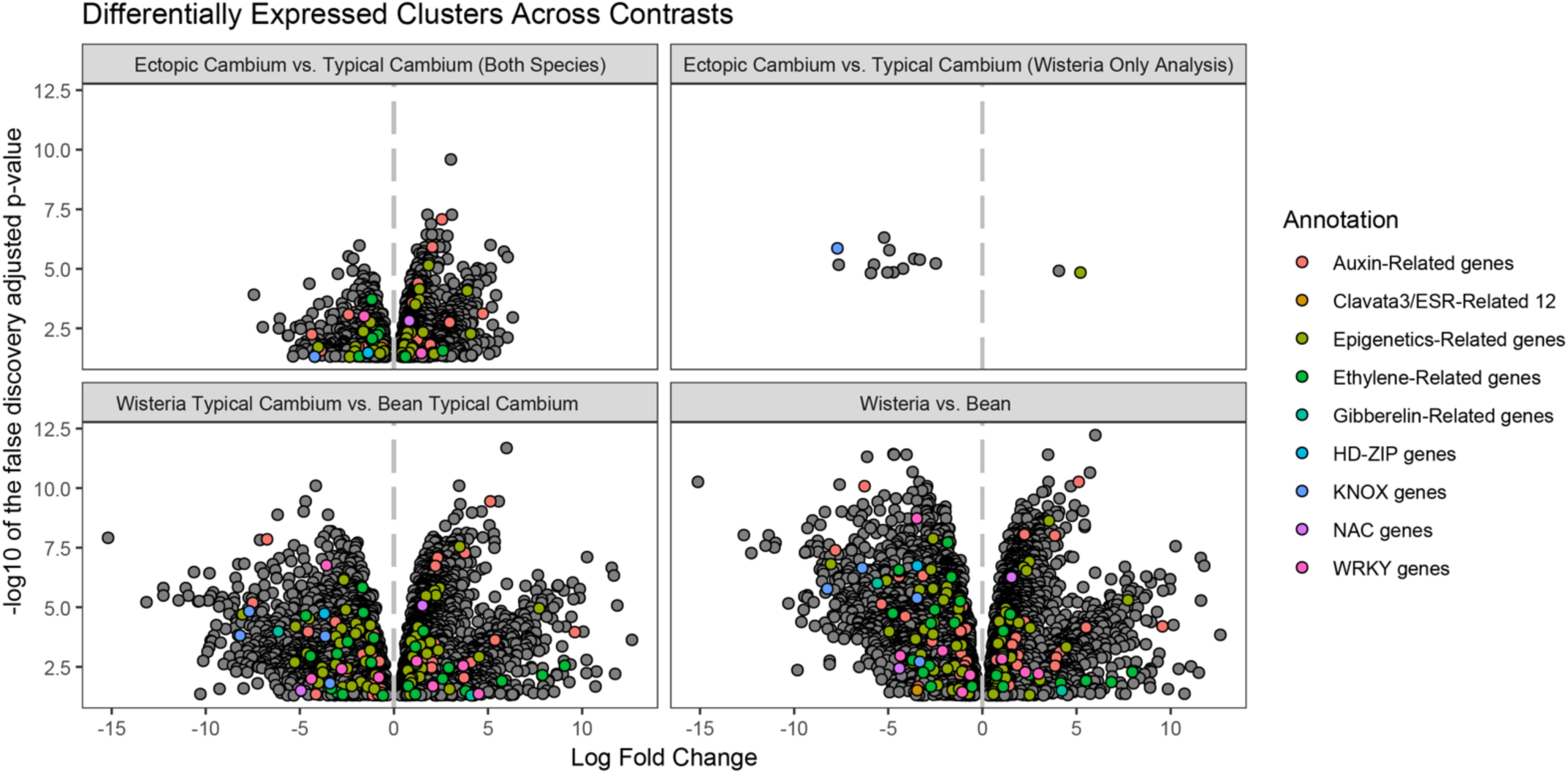
Volcano plot of differentially expressed transcript clusters by contrast. The y-axis is the negative log10 of the false discovery adjusted p-value, while the x-axis is the log fold change. The transcript clusters classified with colors are related to cambium development in other plants and the group annotation is provided in the legend.

**Table 4.** Log fold change (logFC) of differentially expressed transcript clusters in the “Japanese wisteria: Ectopic v. Typical cambia” using dream.

| Cluster/Protein annotation | logFC |
| --- | --- |
| KNOX1 | -7.70 |
| EXLB1_ARATH | -7.63 |
| Cluster_W13 | -5.92 |
| Cluster_W7 | -5.76 |
| Glyco_hydro_18 | -5.22 |
| Bowman-Birk_leg | -5.04 |
| DIOX_N | -4.94 |
| Dimerisation | -4.72 |
| Mem_trans | -4.22 |
| glycine-rich cell wall structural protein 1-like* | -3.62 |
| Aminotran_1_2 | -3.33 |
| Thioredoxin | -2.48 |
| Cluster_W10 | 4.07 |
| Methyltransf_7 | 5.21 |

**Footnote:** (\*) this gene was not originally annotated but retrieved through BLAST analysis after it was identified as significant.

**Table 5.** Log fold change (logFC) of transcript clusters annotated as *KNOX* genes and differentially expressed across contrasts.

| Cluster | Protein annotation* | Japanese wisteria v. Common bean | Typical Cambia: Japanese wisteria v. Common bean | Ectopic Cambia v. Typical Cambia of Japanese wisteria and Common bean | Japanese wisteria: Ectopic v. Typical cambia |
| --- | --- | --- | --- | --- | --- |
| Cluster_W2 | KNOX1 (KNAT2/6) | NA | NA | NA | -7.70 |
| Cluster_9937 | KNOX1 (STM) | -8.24 | -8.17 | -4.22 | NA |
| Cluster_2082 | KNOX1 (STM) | -3.47 | -3.65 | NA | NA |
| Cluster_4072 | KNOX1 (KNATM) | -3.33 | -3.40 | NA | NA |
| Cluster_1332 | KNOX2 (KNAT2/6) | -6.38 | -7.69 | NA | NA |

Footnote: The asterisk (\*) indicates the KNOX protein based on our phylogenetic analysis (see below).

### Identification, characterization, and phylogenetic analysis of KNOX genes analysis

To probe the relationship between *KNOX* genes and EC evolution, we began by testing the hypothesis that there has been a diversification of *KNOX* genes leading to EC. Toward this aim, we broadened our analyses to encompass all nine members of the *KNOX* gene family (based on the *A. thaliana* genome) across 45 seed plant species, including gymnosperms, monocots, and eudicots. Our sampling included species with and without EC (Supplementary Table 5-6). Although retrieved through the tBLASTx search, we removed eight species given their poor alignment, thus potentially erroneous phylogenetic placement (Supplementary Fig. 2).

The final sequence alignment contained 466 sequences (see Supplementary Dataset 1 and 2 for aligned and unaligned nucleotide alignments, respectively). The maximum likelihood phylogeny grouped *KNOX* genes into three main clades that correspond to defined classes (Fig. 3; Supplementary Fig. 1): KNOX1, KNOX2, and KNATM (Fig. 3). In this phylogeny rooted with a representative of *BEL1* genes, KNOX2 diverges first, with the subsequent divergence of KNOX1 and KNATM as sister lineages. The three KNOX classes can be further divided into major clades named according to their putative *A. thaliana* ortholog (Fig. 3; Supplementary Fig. 1). Class KNOX1 includes three main clades corresponding to STM, KNAT1, and KNAT2/6. Class KNOX2 is subdivided into two clades corresponding to KNAT3/4/5 and KNAT7.

**Fig 3.**
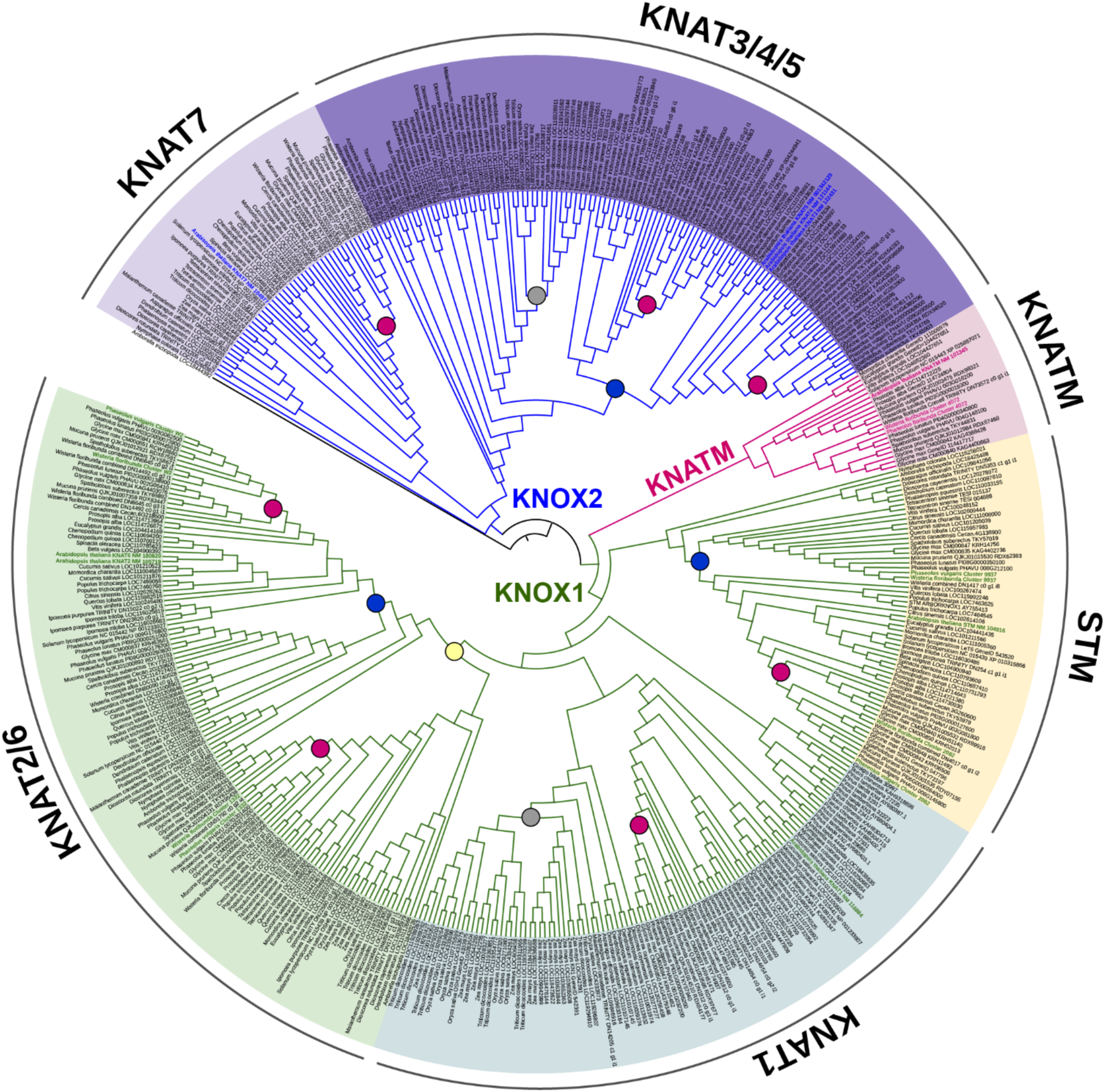
Maximum likelihood phylogenetic tree of KNOX proteins derived from amino acid alignment. The phylogenetic tree includes members of *KNOX* genes obtained from a tBLASTx search, including members of seed plants from 20 families, 33 genera, and 38 species, including 11 species that develop ectopic cambia. KNOX proteins are divided into three classes; KNOX1 (branches in green), KNOX2 (branches in blue), and KNATM (branches colored in magenta). These classes are further divided into clades corresponding to individual genes named after *Arabidopsis thaliana* orthologs, as indicated in bold colors (green, blue, magenta). All recovered transcript clusters annotated for KNOX genes are likewise highlighted in bold colors corresponding to their identified clades (KNOX1, KNOX2, or KNATM). The tree is rooted in a member of the *BEL1* gene family. Circles indicate angiosperm (yellow), eudicots (blue), Poales (grey), and Fabaceae-specific (magenta) duplications.

Using our phylogenetic approach, we were able to determine the orthology of each of the differentially expressed KNOX transcript clusters (see Table 6). All DE clusters belong to class KNOX1: Two of our transcript-clusters group with *A. thaliana* orthologs of STM, one of which is nested within a clade derived from a Fabaceae-specific gene duplication containing the soybean SBH1 reference; two transcript-clusters group with KNAT2/6, one being associated with each of the major clades resulting from gene duplication following the divergence of angiosperms (Supplementary Fig. 1); and one transcript-cluster groups with KNATM (Fig. 3; Supplementary Fig. 1).

The number of KNOX gene copies recovered from each taxon varies as a result of duplication events within major clades representing different KNOX proteins. We recovered duplication events in most lineages with EC including at the base of the angiosperms for KNAT2/6 (indicated by sequences of *Amborella trichopada* in Fig. 3), eudicots for STM, KNAT2/6, and KNAT3/4/5) (indicated by sequences of *Vitis vinifera* in Fig. 3), as well as within the Fabaceae family for all KNOX1 and KNOX2 genes (indicated by sequences of *Wisteria floribunda* or *Spatholobus suberectus* in Fig. 3). Duplication events are also observed in lineages without EC, including the conifers for KNAT1 and the Poales (monocots) for KNAT1 and KNAT3/4/5 (indicated by sequences of *Zea mays or Triticum dicoccoides* in Fig. 3). We did not find an association between the number of *KNOX* genes and the presence/absence of EC across plants (Fig. 4). The *A. thaliana* genome contains nine members of the *KNOX* gene family, but lineages with EC include both increases (e.g., Japanese wisteria with 16 copies and other species of Fabaceae with 17 copies) and decreases (e.g., Amaranthaceae species with five copies), including the possible absence of *KNOX* genes in the EC-producing gymnosperms *Cycas panzhihuaensis* and *Gnetum montanum* (Fig. 4). A Fabaceae-specific duplication in KNATM after the divergence of *Cercis canadensis* is apparent in this dataset (Fig. 4).

**Fig. 4.**
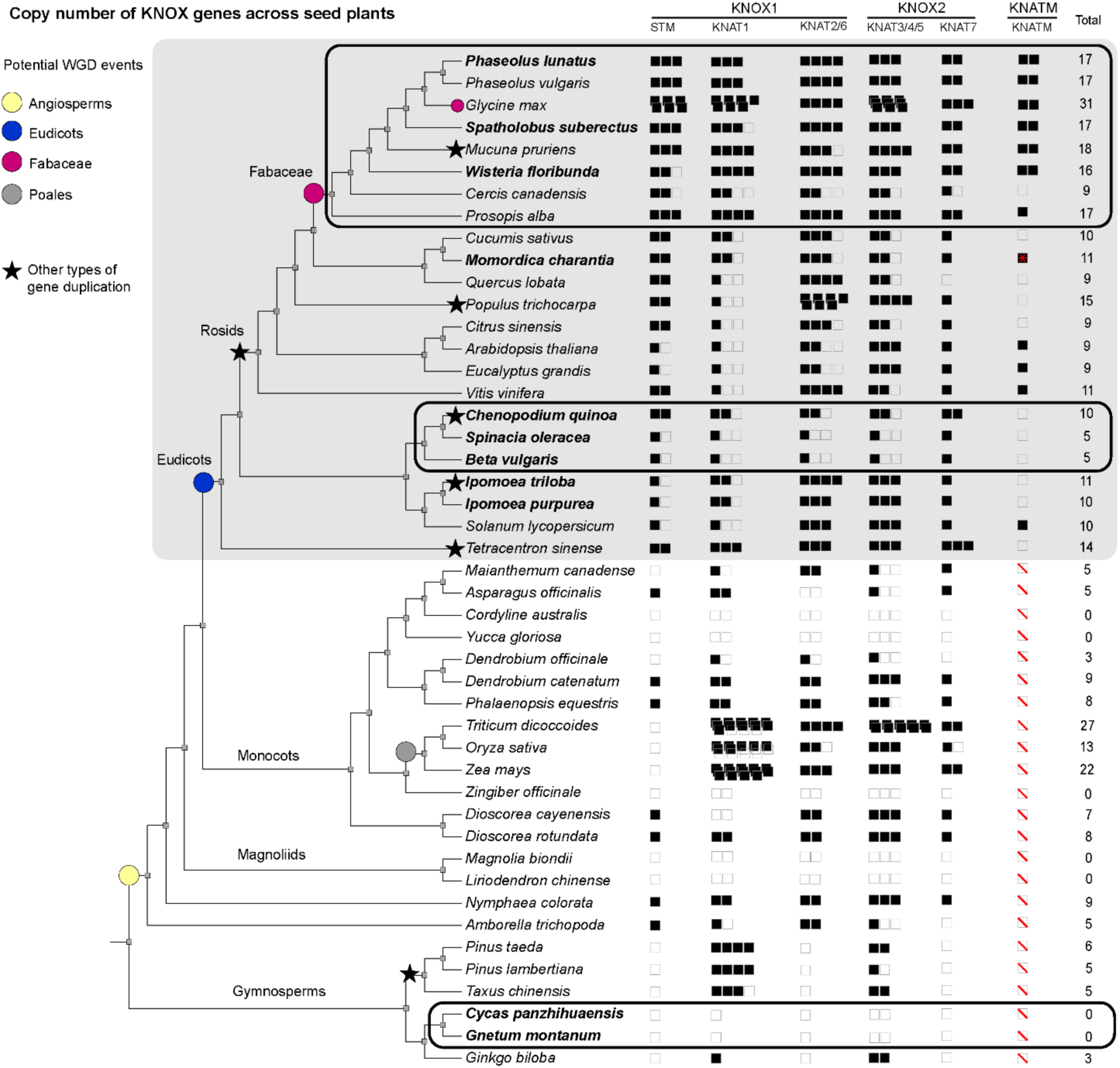
Copy number variation of *KNOX* genes plotted in the species tree from the sampled taxa used in the phylogenetic analyses (Fig. 3). Copy numbers are inferred from the phylogeny in Fig. 3 and associated with duplication events. Squares indicate copy number, with filled squares indicating gene presence, empty squares inferring gene losses, and white crossed squares indicating gene absence. Whole genome duplications (WGD) inferred in Fig. 3 are indicated by colored circles; stars indicate inferred taxon and/or copy-specific duplication events. Expansion of some *KNOX* genes is evident after WGD events in angiosperms (yellow circle), Eudicots (blue circle; grey box), and Fabales (magenta circle). Names of species with ectopic cambia (EC) are in bold. Black ellipses indicate variation in copy number in lineages with EC. All copies were recovered from the blast pipeline to build the phylogeny (Fig. 3), except for *Momordica charantia* KNATM (red asterisk) which was downloaded from Genbank.

**Fig. 5.**
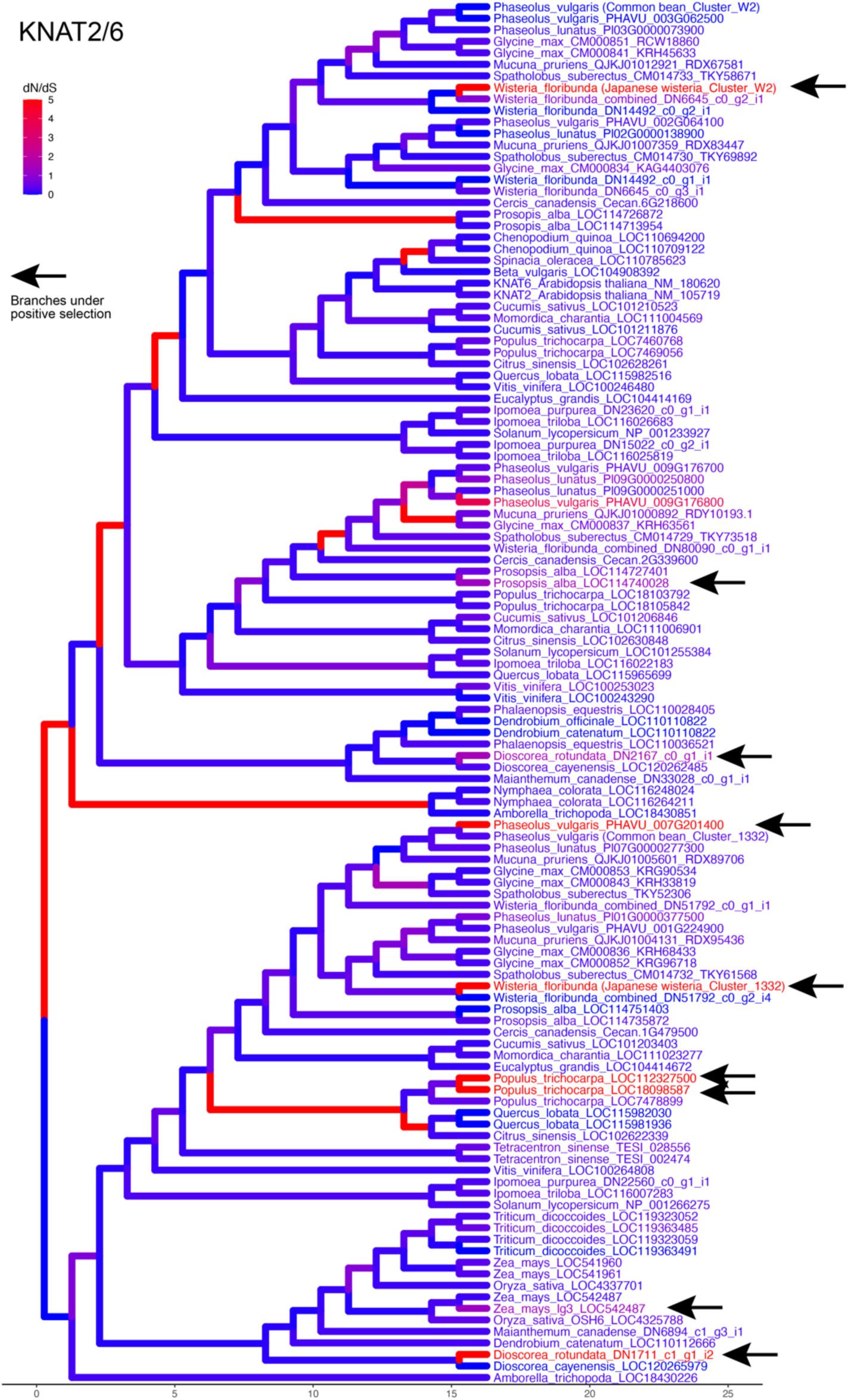
Branch-specific selection on KNAT2/6 gene family. Rooted tree fitted for branch models for KNAT2/6 gene investigated under selection analysis. A unique omega value (dN/dS ratio, ω) is assigned per branch (model 1). Branches in red indicate ω > 1 (positive selection; indicated by arrows), and blue branches indicate ω < 1 (purifying selection).

### Selection analyses

To test whether KNOX genes in lineages with EC are under positive selection (indicative of adaptive evolution) we compared PAML codon-based nested models (models 0 = one omega value for the tree, 1 = a unique omega per branch, 2 = foreground and background omega values^26^). We tested these models in individual gene trees for members of KNOX genes that were differentially expressed in the transcriptome analysis (i.e., STM, KNATM, and KNAT2/6). By comparing foreground (EC present) and background (absent/typical growth) branches (model 2), the only gene set that showed a significant difference was KNAT2/6 (p-value = 0.00406). To detect whether some branches along KNAT2/6 lineages were under positive selection, we employed branch model 1 on the KNAT2/6 phylogenetic rooted tree, which showed nine branches with omega values greater than 1 (ω >1), therefore, indicating positive selection in certain lineages (Supplementary Fig. 3). Out of the nine branches, only two have EC, both of which are sequences differentially expressed from two transcript clusters in the gene expression analysis (*Wisteria_floribunda*_Cluster_W2, ω = 999; *Wisteria_floribunda*_Cluster_1332, ω = 958.234). Taxa under positive selection belonging to species with typical growth include *Dioscorea rotundata* (ω = 999)*, Populus trichocarpa* (ω = 964.749 and 999), and *Z. mays* (ω = 1.29156), as well as some Fabaceae such as *Phaseolus vulgaris* (ω = 999) and *Prosopis alba* (1.27446). Despite statistical tests not supporting differences in synonymous v. nonsynonymous changes for the KNATM protein, results indicate only two branches with higher omega values both of which are species with EC from the Fabaceae including *Phaseolus lunatus* (ω = 1.53725) and *W. floribunda* (ω = 1.15616), representing the KNATM sequence from the DGE analysis (Cluster_4072) (Supplementary Fig. 3). Only one branch belonging to *P. trichocarpa* (ω = 999) is under positive selection in the rooted STM tree (Supplementary Fig. 3).

## Discussion

### *KNOX* genes are likely candidates for modulating EC formation in Japanese wisteria

In Japanese wisteria, ectopic cambia (EC) originate from cortical parenchyma, forming one to several vascular increments that may or may not be continuous^9^. This ontogenetic origin from the cortex is unlike most reported species with EC, where cambia typically form from vascular tissues, e.g., pericycle parenchyma, phloem parenchyma^6^. Until recently, information on how this process occurs at the molecular level was derived solely from experimental studies in *Populus* and *A. thaliana*, whereby mis-expression of conserved genes related to vascular development yielded the formation of EC from the cortex or vascular tissues^6^. Given these data and considering the conserved nature of vascular development across seed plants, we had originally hypothesized only a few developmental pathways likely leading to the formation of EC, i.e., HD-ZIP III and *WOX4* overexpression, similar to the observations from experimental mutants^6^.

Although candidate genes were annotated in the Common bean and Japanese wisteria transcriptome, they were not present in the final analysis of differentially expressed genes due to either not being assigned as one-to-one orthologs across species, not being annotated in one or the other species (e.g., *ANT, AtHB8, WOX4*), not being annotated in both species (e.g., *CLE41/42/44, WOX14*), or being present in both species and not differentially expressed (e.g., *REV*). However, other genes known to be associated with EC formation were not annotated (e.g., *AtHB4*). These results are a combination of differences associated with both biological variation and methodological limitations. Here we found evidence that members of the *KNOTTED1-like homeobox (KNOX)* gene family are among the top candidates for genes related to the formation of EC. Our KNOX phylogeny recovered three KNOX classes with relationships among the classes similar to those recovered in previous studies^27^, while the phylogenetic position of subclasses (e.g., KNOX1) in some lineages (e.g., conifers) differ across different studies, including our tree^28,29^. Placing our five differentially expressed KNOX transcript clusters into a phylogenetic context, we show that all copies fall into either KNOX class I or KNATM. We will focus our discussion on these two classes.

KNOX1 genes have been reported to act as major regulators of organ initiation, tissue proliferation, and meristem maintenance, including the formation of vascular cambium^19,30–32^. In *Populus*, the overexpression of ARK1 (ortholog of *SHOOTMERISTEMLESS* (*STM*) in *A. thaliana*) generated dispersed/unequal differentiation of cambium derivatives during the secondary growth^30^, while the overexpression of ARK2 (ortholog of KNAT1 in *A. thaliana*) promoted a wider cambial zone^31^. In *A. thaliana*, KNAT1 mutants exhibited reduced radial growth although meristematic activity was maintained, which indicated functional redundancy with other genes (e.g., KNAT6) in the transcriptional regulation of cambial activity and differentiation of vascular tissues^19,33,34^. In Japanese wisteria, we saw downregulation of a transcript-cluster identified as KNAT2/6 in the “Japanese wisteria: Ectopic v. Typical cambia” contrast (Table 3). Since KNAT6 is known to be expressed in the vascular cambium and vascular tissues of most species^33–35^, although absent in the cambial zone of some Cactaceae^36^, the misexpression of these genes in the “Japanese wisteria: Ectopic v. Typical cambia” contrast may suggest their involvement in EC development. Other evidence of the involvement of KNAT6 in vascular oddities may be noted from the higher transcript levels of KNAT6 in *knat1* and *pny* (*PENNYWISE* also known as *BEL1-LIKE HOMEODOMAIN 9*) *A. thaliana* mutants. These plants show vascular defects ranging from increased cambial activity between the phloem and xylem cells^37^ to extravascular bundles that enlarge to reduce interfascicular space^38^. Curiously, StPOTH1, an ortholog of KNAT6 in *Solanum tuberosum* L., may also be involved in the development of potato tubers^39^, which have been described as having atypical vascular tissue formation^40^. Nevertheless, just as the mechanism by which KNAT6 functions in vascular development remains largely unclear in model species^30,33,34,38^, additional studies will be required to clarify how KNAT2/6 plays a role in EC development, as well as the involvement of other KNOX proteins (e.g., STM), which is also downregulated in the contrast comparing “Ectopic Cambia v. Typical Cambia of Japanese wisteria and Common bean”. The functional validation of KNOX proteins in relation to vascular development should address gene expression dynamics and potential interactions, which suggest that they may share similar functions^41^ as well as act in antagonistic relationships (e.g., KNAT1 and KNAT6)^34,42^, including in other systems such as leaf development^43^.

In the cambium regulatory network of woody plants, KNOX proteins (e.g., KNAT1 and STM) are downstream of the *TDIF-PXY-WOX4* signaling pathway, a conserved module responsible for cambium development in woody plants^1,13,14^. These observations suggest that EC development in Japanese wisteria probably derives from modifications to the *TDIF-PXY-WOX4*, perhaps leading to changes in the regulation of downstream *KNOX* genes. That is also evidenced by the findings that KNAT1 was downregulated in *wox4 A. thaliana* mutants with cambia formed ectopically^19^ and by the fact that the overexpression of *WOX4* alone resulted in EC in *Populus*^18^. In addition, *knat1 wox4* mutants exhibited severe defects in cambium proliferation in *A. thaliana,* however, cambium initiation and activity were still detected, indicating redundancy among cambium regulators^19^. The manipulation of these transcription factors (e.g., *WOX4*) that are putative upstream regulators of KNOX1 genes (e.g., KNAT1) provides evidence of their involvement in vascular development and potentially EC formation. Future studies investigating the expression of these KNOX proteins in EC formation will shed more light on their role in vascular development and the development and evolution of EC in natural systems.

### Both constitutively and injury-induced ectopic cambia converge on similar molecular processes

In this study, we found downregulation of KNOX proteins within the constitutively formed EC of Japanese wisteria. *KNOX* genes also emerge as important regulators underpinning the ability of perturbed *Populus* trees to form de novo cambia post-girdling^20^ (see Supplementary Table 2 and Supplementary Methods for a comparison of cambial markers across papers on vascular development). In girdled *Populus*, KNOX genes (i.e., KNAT1, KNAT3, KNAT6) are up-regulated in most stages (e.g., dedifferentiating xylem parenchyma, differentiating cambium). However, they become down-regulated in the final stage when a regenerated cambium is finally observed 12 days after girdling (Supplementary Table 2). KNOX genes are also down-regulated in the cambial zone of unperturbed *Populus*^22^ (i.e., KNAT1, KNAT2, KNAT6; Supplementary Table 2), corroborating their reduced expression profiles in mature stages of cambium development. Conversely, *KNOX* genes are up-regulated in mutants in which ectopically-formed cambia are observed, such as *Populus HB4*-repressed (i.e., *KNAT3*^17^) and the *A. thaliana* w*ox4* (i.e., KNAT1^19^) (Supplementary Table 2). Thus, *KNOX* genes appear to have been independently co-opted in the molecular regulation of de novo cambia in both constitutively formed (*Wisteria* vines) and injury-induced (girdled stems of *Populus*) conditions.

In addition to *KNOX* genes, other major cambium regulators such as phytohormones are differentially expressed in constitutively and injury-induced EC (see Supplementary Table 3 for gene families differentially expressed across different studies). Previous investigations suggest that *ETHYLENE RESPONSE FACTORS* (*AP2/ERF*) may be upregulated in the absence of *PXY*, defining an alternative pathway to the *PXY/CLE41* module that regulates cell division in vascular tissue development^44^. Notably, *PXY/CLE41* (as well as *TDIF* and *WOX4* which may also be part of this signaling pathway) are not retained as differentially expressed genes in the comparisons among cambium types in our study, while *AP2/ERF* genes are differentially expressed in both Japanese wisteria and girdled *Populus*, along with other genes from the ethylene signaling cascade (e.g., *EIN2* and *EIN3*) (Supplementary Table 3). Auxin-related genes are considered master regulators of vascular development because they induce the expression of other conserved transcription factors (e.g., *WOX4*, *PXY,* and *AtHB8*)^45,46^. In line with those findings, changes in gene expression patterns in auxin-related genes in both injury-^21,47^ and non-injury-induced systems (this study) add evidence to shared mechanisms in de novo cambium formation (e.g., *ARF, AUX, PIN6, TIR;* Supplementary Table 3). The re-establishment of typical auxin pathways in cambium regeneration in injury-induced cambia is consistent with typical cambium development, and auxin is also sufficient to induce cambium regeneration *in vitro*^47^. Investigations into the role of auxin in EC initiation are a required and exciting venue for future research on vascular biology.

Lastly, constitutively and injury-induced EC share signatures of epigenetic regulation. We found that numerous methyltransferases were differentially expressed across cambium types, being upregulated in Japanese wisteria EC (Supplementary Tables 1, 3). EC in lianas such as Japanese wisteria are potentially triggered by external environmental factors such as mechanical and hydraulic constraints linked to their climbing habit. Findings from *Bignonia magnifica* W.Bull show that adding external support for these vines to climb triggers changes in the xylem anatomy from self-supporting to liana-like with associated changes in the transcription profiling of the cambium^48^; thus external cues relate to anatomical changes in these vines (see Supplementary Table 4 for comparison of differentially expressed genes among woody vines; see also Supplementary Methods for further explanation). Although the epigenetic regulation of the vascular cambium remains poorly understood, recent evidence suggests a mechanism involving histone acetylation at the *WOX4* promoter^49^. The association between external factors, gene regulation, and atypical vascular architectures has also been reported in swallow underground storage organs, which also thicken through parenchyma proliferation, additional vascular bundles, or EC itself (e.g., beets)^40^. A notable similarity across underground storage organs and EC in Japanese wisteria is the role of expansin encoding genes (associated with cell wall loosening), in cell division and proliferation. We found expansin downregulated in the Japanese wisteria EC, which was also reflected in the underground storage organs of *Bomarea multiflora* (L.f.) Mirb.^50^. However, their specific role is yet to be further elucidated given that these genes are up-regulated or down-regulated in different vascular systems (e.g., types of underground storage organs or EC)^40^.

The enriched GO classes of genes that were differentially expressed in the “Ectopic Cambia v. Typical Cambia of Japanese wisteria and Common bean” contrast (Table 3) include those related to mRNA processing, splicing, and binding, as well as nucleus, nucleolus, and polymerase activity. These terms may be associated with cambium formation as chromatin alterations, increased transcription, and histone modification are necessary for cell dedifferentiation and cell cycle re-entry during plant regeneration^51–53^, as also indicated for the injury-induced EC development in *Populus*^20^. This molecular program may also explain how cortical and pericyclic cells are induced to undergo cell division independently of stem cell lineage in regeneration systems^51,54^. In addition, previous research shows that different levels of mRNA processing factors involved in pre-mRNA splicing are critical for plant development, including the differentiation process before initiation of cell division^55^. Taken together, these observations indicate that similar molecular events may be involved in cambial cell competency during EC development.

According to our KNOX phylogeny, gene copy number is variable across lineages, but there is no association with EC evolution. There are instances of higher copy number of *KNOX* genes in lineages with EC (e.g., Fabaceae species), while in other lineages both an increase and a reduction are observed (e.g., Amaranthaceae) in comparison to the nine copies of *A. thaliana*. Copy number variation is related to duplication events identified at various taxonomic levels. Fabaceae-specific duplications were observed for most genes set within the KNOX1 and KNOX2 classes, as well as within KNATM*—*which has been reported only in a few eudicot lineages^56^. Our study suggests that the expansion of *KNOX* genes is derived from either whole genome duplications (WGD), segmental, or tandem duplications but the specific underlying cause for each has not been identified in this study. For instance, both WGD or segmental duplications in multiple genes could explain an increase in copy number in eudicots or the Fabaceae.

Polyploidy events are well-known in the evolution of the genus *Glycine*^57^ and the studied grasses^58,59^, which is more likely to explain higher copy numbers, although segmental and tandem duplications have also been reported as the mechanisms in the expansion of *KNOX* genes in grasses (e.g., *Triticum aestivum*^60^). In addition, paleopolyploidization events have been reported for all sampled angiosperm genera/species, however, most have gone through diploidization events (see Supplementary Table 7 for a list of polyploidization events) indicating that associating copy number variation with WGD should be taken with caution. Differences in copy number within some lineages may also be a result of sampling transcriptomes rather than genomes for some species (e.g., *Maianthemum canadense, Ipomoea purpurea*).

Our selection analyses indicate that KNOX genes (e.g., KNAT2/6) are under positive selection, including two sequences of Japanese wisteria from the transcript clusters obtained in the differential expression analysis. In addition to Japanese wisteria, however, contigs from other species with or without any vascular oddity are also under positive selection. Cases such as *Dioscorea,* which forms underground storage organs and potentially atypical vascular tissue formation, may corroborate selective pressures in the evolution of copies of KNAT2/6 associated with vascular oddities. However, we found a positive selection of KNAT2/6 in species without any vascular aberration (e.g., *Populus*), thus further analyses are necessary to investigate the gene-phenotype relationship.

In conclusion, our work provides anatomical and molecular insights into the unusual emergence of EC formation from cortical parenchyma in Japanese wisteria vines. Amongst the top most differentially expressed genes are numerous conserved genes involved in hormone signaling, cell division and differentiation, and epigenetic regulation. Our findings reveal *KNOX* genes as important factors modulating EC development in Japanese wisteria. Although none of the HD-ZIP III genes (i.e., *REV, AtHB7, AtHB4*) generating EC in experimental mutants were retained as a differentially expressed gene in Japanese wisteria, we cannot rule out that this pathway represents an alternative route for the emergence of de novo cambia, since EC have arisen through the misexpression of HD-ZIP III genes in experimental mutants. Future investigations leveraging spatial, and time-specific transcriptomes will further elucidate the genetic mechanisms controlling EC and its similarities with de novo cambia in other natural and induced vascular phenomena in plants (e.g., cambium regeneration, tuber formation). This knowledge will inform the contribution of conserved genetic pathways to the repeated evolution of the climbing habit in plants.

## Methods

### Plants and stem sample collection

*Wisteria floribunda* (Japanese wisteria) plants were collected in two populations growing in Ithaca, NY, United States. We performed DNA barcoding using the chloroplast intergenic spacer gene “ndhJ-trnF” to confirm the identification of plants belonging to the species *W. floribunda*^10^*. Phaseolus vulgaris* (genotype L88-57; Common bean) plants were grown at Cornell University Agricultural Experiment Station Guterman greenhouses (Ithaca, NY) under the following conditions: 14h light, 75 F daytime temperature, 65 F nighttime temperature, and ambient humidity. Samples from these plants were used for developmental anatomy and RNAseq analyses (see methods below).

### Histology and microscopy

Stem samples were hand-sectioned (with the aid of a razor blade) and processed to generate temporary slides or processed using standardized methods in wood anatomy (i.e., embedded in polyethylene glycol, sectioned, and stained) to generate permanent slides^61^. Slides were imaged using a light microscope (Olympus BH2) with an attached digital camera (AmScope MU1000).

### RNA sequencing and data processing

#### Sampling and RNA isolation

Developed stem samples were cut into small blocks containing cambium, secondary xylem, and secondary phloem, which were sectioned (∼12 mm thick) using a cryostat at -20°C (tangential cryosectioning approach^22^) to obtain the tissue for total RNA isolation with RNeasy Mini Kit (QIAGEN). For RNA extraction, about 100 sections of each sample containing cambium, secondary xylem, and secondary phloem were used for each replicate. Six biological replicates were used for the typical cambium of Common bean and ectopic cambia (EC) of Japanese wisteria, while five replicates were used for the typical cambium of Common bean. The same adult plants used for the comparative transcriptomes were used for anatomical studies (as indicated above).

#### RNA integrity and quantification

Total RNA was quantified using RNA Assay Kit in Qubit® 2.0 Fluorometer (Life Technologies, CA, USA), and RNA integrity was assessed using an AATI Fragment Analyzer (Agilent Technologies, CA, USA). Purification, library preparation, and Illumina sequencing were performed by Novogene Co., Ltd (Davis, California). Quality control was performed from the RNA samples to the final data using Qubit® 2.0, and RNA integrity was assessed using the RNA Nano 6000 Assay Kit of the Agilent Bioanalyzer 2100 system (Agilent Technologies, CA, USA).

#### Library preparation for Transcriptome Sequencing

RNA library for poly(A)-dependent RNA-seq was prepared using NEBNext Ultra II RNA Library Prep Kit for Illumina (New England BioLabs). The libraries were subjected to 150 bp paired-end sequencing using NovaSeq 6000 with the v1.5 S4 reagent kit. The library was checked with Qubit and real-time PCR for quantification and bioanalyzer for size distribution detection.

#### Transcriptome Assembly and Annotation

Raw paired-end reads were trimmed for adapter sequences using an overlap stringency of three, a Phred score cut off of twenty, and a minimum sequence length of 50 base pairs (bp) using Trim Galore! version 0.6.6^62^. De novo transcriptome assemblies were developed for *P. vulgaris* and *W. floribunda*. For each trimmed dataset, the range of potential kmer values was estimated with Kmergenie version 1.7016^63^. De novo assemblies were generated following part of the TransPi pipeline that uses a multi-assembler approach followed by merging assemblies, reducing complexity, and annotation^36^ (see Supplementary methods section “TransPi pipeline” for additional details). Within the TransPi pipeline, the UniProt database for Fabales (UniProt Taxon ID 72025^24^)was used for annotation (*see Supplementary methods section* “*Limitations of the one-to-one orthologs approach”* for additional details).

#### Differential Expression and Gene Ontology Enrichment

Trimmed reads were subset to 20 million reads, mapped to their respective transcriptomes with Hisat2 version 2.2.1^64^ including all multiple alignments, and the mate-pair information of the resulting SAM files were fixed using Picard Tools version 2.27.5 FixMateInformation before being converted to BAM files with SAMtools version 1.14^65^. Corset version 1.07^66^ was used to group the mapped transcripts into transcript clusters and estimate read counts for each cluster for both transcriptomes separately. Each cluster was annotated using the Trinotate annotation of the longest sequence within the cluster. Clusters were retained if at least two samples in each species had 0.5 counts per million. For the cross-species comparison, Orthofinder2 version 2.5.2^67^ was used with our protein annotations from TransDecoder, and the proteomes of *Vigna unguiculata* (UniProt Taxon ID: 3917), *Cajanus cajan* (UniProt Taxon ID: 3821), *Lotus japonicus*^68^, *Medicago truncatula*^69^ and *Mucuna pruriens* (UniProt Taxon ID: 157652) from UniProt^24^ and Phytozome^70^ to identify orthologs between *P. vulgaris* and *W. floribunda*. We identified one-to-one orthologs at the transcript cluster level by removing any clusters with more than one ortholog group and any ortholog groups that contained sequences from more than one cluster. Differential cluster expression were tested using limma^71^ and voom^72^ comparing between species regardless of cambium type, between cambium types regardless of species, between the typical cambium of each species, and between the ectopic and typical cambium of Japanese wisteria (*see Supplementary methods section* “*Limitations of the one-to-one orthologs approach”* for additional details). Gene ontology enrichment was tested using the differentially expression transcript clusters for each using GOseq^73^. To better leverage the repeated sampling within an individual, Japanese wisteria clusters were tested separately from the cross-species comparison using <u>d</u>ifferential expression for <u>re</u>pe<u>a</u>ted <u>m</u>easures (dream^25^) with voom^72^ and variancePartition^74^ to compare across cambium types with individual as a random effect. Gene ontology enrichment for differentially expressed clusters was implemented in GOseq. For comparison with the cross-species analysis, ortholog groups were subsequently mapped back to the differentially expressed clusters identified by dream.

### Phylogenetic and Selection analyses

To further investigate the role of *KNOX* genes in EC formation across seed plants, we performed a phylogenetic analysis and selection analysis for this gene family. Candidate genes were identified from available genomes and transcriptomes using a BLAST pipeline that was previously used for investigating gene evolution across the angiosperms^75^. Coding sequences (cds) were downloaded from Genbank or TreeGenes for 39 species and raw RNA-seq files were downloaded from the SRA for six species (Supplementary Table 5). In total, 45 species were included in the analysis, 11 containing EC. Raw sequencing files were cleaned with fastp version 0.23.3^76^ and assembled with Trinity version 2.14.0^77^ with the following flags: --max_memory 200G --no_normalize_reads --min_contig_length 250 --trimmomatic.

Reference sequences of candidate genes from *A. thaliana* were downloaded from Genbank and a tBLASTx search was performed using blast version 2.13^78^. A maximum of 30 hits per taxon were retained if the matching length was 10 bp and an e-value of 0.0001. A fasta file containing all retrieved sequences was generated and checked for duplicates using the sRNAtoolbox^79^. Twenty-one sequences from different *KNOX* genes and taxa were downloaded from Genbank and included as additional references to identify specific clades (Supplementary Table 6). A species tree to depict this sampling (Fig. 4) was built using the R package V.PhyloMaker2 using scenario 3 for taxa binding^80^. Finally, the sequences of five transcript clusters identified as KNOX genes were included in the phylogenetic analysis. The resulting fasta file for phylogenetic reconstruction contained 466 sequences. These contigs were first aligned with MAFFT version 7.52^81^ with the following options: --auto --adjustdirectionaccurately --op 3 --leavegappyregion. The sequences were then realigned with the codon-aware aligner MACSE version 2.07^82^ and the alignSequences command. The resulting alignment was cleaned up with trimAL version 1.4.1^83^ with the - gappyout option before converting premature stop codons to NNN while retaining the final stop codons with the exportAlignment command in MACSE, which were cleaned up with trimAL version 1.4.1^83^. Tree building was performed reiteratively to exclude poorly aligned and gappy contigs (see Supplementary Methods). Trees were generated with RaxML version 8.2.12 with the GTRGAMMA model of molecular evolution and 100 bootstrap replicates^84^.

For further investigation of specific candidate genes, a fasta file containing the nucleotide sequences of specific *KNOX* genes identified in the previous tree was aligned with MAFFT version 7.52, realigned with the codon-aware aligner MACSE version 2.07, and then cleaned with trimAL version 1.4.1. The inferred phylogeny and cleaned alignment were inputs into PAML version 10.6^26^ for codeml analysis to explore signatures of selections. Both model = 0 (one omega value for the entire alignment), model = 1 (omega value for each branch), and model = 2 (foreground and background omega values) were conducted on both the rooted and unrooted trees. Branch-specific omega values were plotted on the phylogeny with the R package ggtree version 3.6.2^85^.

## Reporting summary

Further information on research design is available in the Nature Portfolio Reporting Summary linked to this article.

## Data availability

The datasets generated and/or analyzed during the current study are publicly available. The RNA sequencing data is available in SRA (SRR28389658, SRR29802342-SRR29802358). Other datasets generated in this study have been deposited in the GitHub database (https://github.com/anthonysnead/Fabaceae-Ectopic_Cambia_Transcriptomics).

## Code availability

A custom R script was written for gene expression analyses and is available on GitHub (https://github.com/anthonysnead/Fabaceae-Ectopic_Cambia_Transcriptomics).

## Acknowledgments

We thank undergraduate research technician Danielle C. Sonnenleiter (Cornell University) for assistance in data collection, past and current members of the Onyenedum lab for continuous feedback, especially Angelique A. Acevedo for assistance in growing the bean plants, and Mariane S. S. Baena for insightful discussions. We also thank Jocelyn Rose’s lab at Cornell University for allowing access to the cryostat. We would like to acknowledge the Arnold Arboretum of Harvard University for providing access to the living collections and financial support through a Sargent Award for Visiting Scholars (I.L.C.N.). This work was supported in part through the NYU IT High Performance Computing resources, services, and staff expertise as well as the Boyce Thompson Institute’s Computational Biology Center and Cornell’s BioHPC. This work was funded by startup laboratory funds from Cornell University’s College of Agriculture and Life Sciences and New York University, and NSF 2237046 to J.G.O.

## Author contributions

Conceptualization: I.L.C.N., J.G.O. Methodology: I.L.C.N., J.B.L., C.D.S, A.A.S., J.G.O. Investigation: I.L.C.N., J.B.L., A.A.S. Data curation: I.L.C.N., A.A.S Writing – Original Draft: I.L.C.N., J.B.L., A.A.S., J.G.O. Writing – Review & Editing: I.L.C.N., J.B.L., A.A.S., C.D.S., J.G.O.

## Additional information (Supplementary information)

**The online version contains supplementary material available at:** https://github.com/anthonysnead/Fabaceae-Ectopic_Cambia_Transcriptomics

**Supplementary Table 1**: All significantly differentially expressed transcript clusters across contrasts.

**Supplementary Table 2**: Comparison between differentially expressed cambial markers across seven studies.

**Supplementary Table 3**: Comparison between differentially expressed genes in this study and induced cambium generation in *Populus*.

**Supplementary Table 4**: Comparison between differentially expressed genes from transcriptomes of woody vines.

**Supplementary Table 5**: List of taxa and source of data used for molecular phylogenetic analysis.

**Supplementary Table 6**: List of taxa downloaded from GenBank used as additional references for molecular phylogenetic analysis.

**Supplementary Table 7**: Number of paleopolyploid events in each species in this study.

**Supplementary Figure 1:** Schematic phylogram of the *KNOX* gene family indicating the three classes, respective genes, and duplication events.

**Supplementary Figure 2:** Schematic phylogram of the *KNOX* gene family including deleted species.

**Supplementary Figure 3:** Rooted tree fitted for branch models for the three genes investigated under selection analysis.

**Supplementary Dataset 1**: Nucleotide alignment for 466 KNOX sequences used in this study.

**Supplementary Dataset 2:** Unaligned fasta file for 466 KNOX sequences used in this study.

**Supplementary Methods**: Description of how Supplementary Tables 2-4 were developed, steps for the KNOX phylogeny, and details on the transPi pipeline.

## Supplementary Figures (legends)

**Fig. S1. Schematic phylogeny of the *KNOX* gene family indicating the three classes with respective genes.** Numbers indicate bootstrap value. Branches of major clades are color-coded as follows: eudicots (green), monocots (dark red), *Amborella* +*Nymphaea* (black), and gymnosperms (yellow). Black arrows indicate the position of *Arabidopsis thaliana* references and names of additional references included in the phylogeny are shown in light green (see also Table S4 and S5). The sequences differentially expressed in our study are shown in pink (indicated with arrows). Circles indicated potential whole genome duplications according to Fig. 3 and Fig. 4.

**Fig. S2.** Schematic phylogram of the *KNOX* gene family including deleted species. The nine species indicated in the purple clade were not retained for the final KNOX phylogeny.

**Fig. S3**: Rooted tree fitted for branch models for the three genes investigated under selection analysis. The arrows indicate the branches under positive selection, *i.e.*, omega values greater than 1 (ω >1). **a** KNATM. **b** STM. Arrows indicate branches under positive selection.

## Supplementary Methods S1

### Building Supplementary Tables S2-S4

To construct **Supplementary Table S2**, we utilized two gene sets as queries across referenced papers. The first set comprises nine *KNOX* genes identified in *Arabidopsis thaliana*, also used in phylogenetic analysis (see Methods). The second set includes 20 genes enriched in single-cell transcriptome analysis of vascular meristems in *Populus*^1^. Gene names (acronyms in column C of Supplementary Table 2) were obtained by querying the NCBI database for gene names (column D) and loci (column E). Column F information was sourced from the Gene Ontology database (https://amigo.geneontology.org/) using *A. thaliana* locus IDs (column E), emphasizing ontology descriptions linked to ectopic cambia (EC) development. Original annotations are in Supplementary Table 1 and referenced studies.

Selected genes (by locus number or acronyms in column C) were cross-referenced with differential gene expression databases from our study and seven others focusing on cambium in *A. thaliana* and *Populus*. These papers fall into two groups: (1) studies without mutant lines such as the wild *Populus* cambium transcriptome^2^ and cambium regeneration after girdling^3^, detailed further in **Supplementary Table S3** due to relevance to EC. (2) Studies with mutant lines including KNAT1 (=ARK2) in *Populus* and *A. thaliana* mutants reported with EC^4–8^. For studies with multiple mutants, each line’s gene expression is shown in separate columns. Studies without differential gene expression data were not included^9^.

In **Supplementary Table S2**, cells are labeled “down-regulated” or “up-regulated” based on log fold change (logFc) from original studies, except for one study which is represented by its “rC/diC” comparisons^3^. Other studies indicated log-fold change through plots^2^ and those were interpreted accordingly.

**Supplementary Table S4.** was created using differentially expressed genes listed across the sole studies on liana stem transcriptomes^10,11^, and cross-checked with our study.

### Summary of steps to generate KNOX phylogeny

(Files are available on https://github.com/anthonysnead/Fabaceae-Ectopic_Cambia_Transcriptomics).

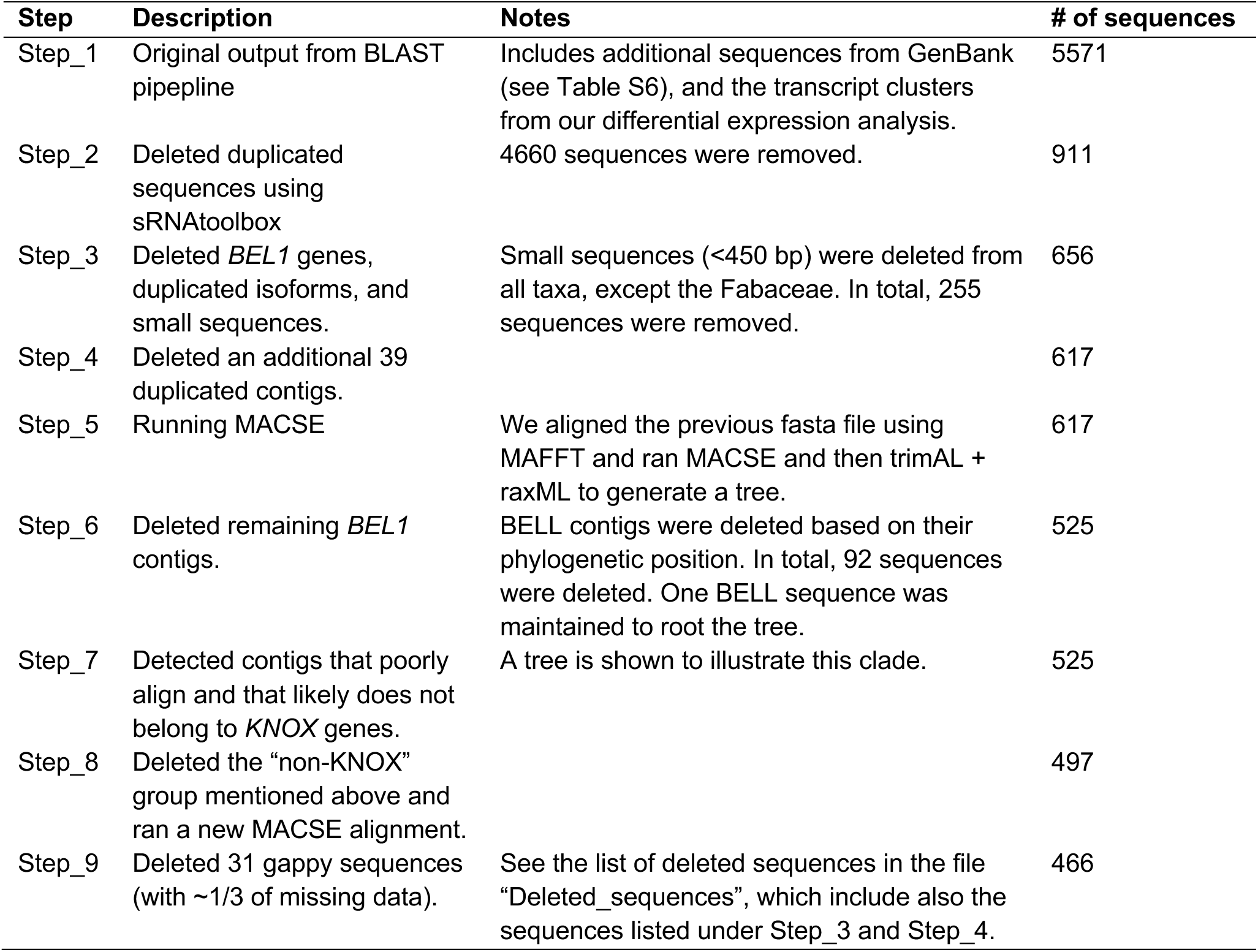

| Step | Description | Notes | # of sequences |
| --- | --- | --- | --- |
| Step_1 | Original output from BLAST pipeline | Includes additional sequences from GenBank (see Table S6), and the transcript clusters from our differential expression analysis. | 5571 |
| Step_2 | Deleted duplicated sequences using sRNAtoolbox | 4660 sequences were removed. | 911 |
| Step_3 | Deleted *BEL1* genes, duplicated isoforms, and small sequences. | Small sequences (<450 bp) were deleted from all taxa, except the Fabaceae. In total, 255 sequences were removed. | 656 |
| Step_4 | Deleted an additional 39 duplicated contigs. |  | 617 |
| Step_5 | Running MACSE | We aligned the previous fasta file using MAFFT and ran MACSE and then trimAL + raxML to generate a tree. | 617 |
| Step_6 | Deleted remaining *BEL1* contigs. | BELL contigs were deleted based on their phylogenetic position. In total, 92 sequences were deleted. One BELL sequence was maintained to root the tree. | 525 |
| Step_7 | Detected contigs that poorly align and that likely does not belong to *KNOX* genes. | A tree is shown to illustrate this clade. | 525 |
| Step_8 | Deleted the “non-KNOX” group mentioned above and ran a new MACSE alignment. |  | 497 |
| Step_9 | Deleted 31 gappy sequences (with ~1/3 of missing data). | See the list of deleted sequences in the file “Deleted_sequences”, which include also the sequences listed under Step_3 and Step_4. | 466 |

### TransPi Pipeline

All de novo transcriptomes were constructed with the TransPi pipeline^12^. Briefly, assemblies were generated for each dataset using rnaSPAdes version 3.14^13^, SOAPdenovo-Trans version 1.03^14^, TRANS-ABYSS version 2.0.1^15^, Velvet version 1.2.12/Oasis version 0.2.09^16,17^ for kmer values 25, 35, 45, 55, and 65 along with Trinity version 2.9.1 ^18^ before being merged and reduced with the EvidentialGene pipeline version 2019.05.14^19^. The EvidentialGene pipeline reduces the complexity of the transcriptomes by collapsing duplicates, clustering protein sequences, and performing similarity searches between the transcripts with a combination of CD-HIT^20^ and BLAST version 2.2.31^21^. Quality assessment was done using TransRate version 1.0.3 (Smith-Unna et al. 2016) for assembly statistics, Hisat2 version 2.2.1^22^ to assess mapping efficiency and <u>B</u>enchmarking <u>U</u>niversal <u>S</u>ingle-<u>C</u>opy <u>O</u>rthologs (BUSCO) version 5.5.0cv1 to evaluate the completeness of expected universal single copy genes with the fables database^23^. For annotation, TransDecoder version 5.5.0^24^ was employed to denote open reading frames (ORFs) and complete homology searches of the ORFs against proteins via BLAST to preserve ORFS with potential functional significance). HMMER version 3.3^25^ was used to complete protein domain searches against the PFAM database. Trinotate version 3.2.0^26^ was used for functional annotation and DIAMOND version 0.9.30^27^ was used for the similarity searches between the transcripts and the UniProt database for fables (UniProt Taxon ID 72025^28^) The final annotation report from Trinotate included the evolutionary genealogy of genes: Non-supervised Orthologous Groups (eggNOG), Kyoto Encyclopedia of Genes and Genomes (KEGG), and Gene Ontology (GO) as well as protein alignments.

### Limitations of the one-to-one orthologs approach

It is important to note that a limitation of working with transcriptomes instead of genomes is that our clusters may not accurately represent genes, and we have been careful to refer to these as transcript clusters to reflect this uncertainty. Genes may have orthologs across species, but the corset clustering might split a real gene into two clusters, affecting our ability to identify one-to-one orthologs. It is also important to note that not all genes are likely annotated in the transcriptomes. Because RNA-seq investigates relative gene expression, reads were subset to a standard 20 million reads before quantification which could reduce the counts of rare genes. Additionally, when filtering for lowly expressed genes by biological treatment, a threshold (i.e., 10 counts) is set for the number of reads that need to be aligned to a given transcript cluster for at least two replicates for it to be retained for downstream analysis. Therefore, if at least two samples in the species do not have at least 10 reads aligned to that transcript cluster, it is discarded. Generally, these are considered genes with extremely low expression and do not contain enough information to be statistically valid. Therefore, three main scenarios may lead to conserved genes not being represented in the final analysis: (1) not assigning one-to-one orthologs due to gene splitting across clusters, (2) unsuccessful protein annotations, and (3) no differential expression in the current analysis, which may be affected by filtering.

