## Supplementary figures and images for "Ectopic cambia in Japanese wisteria (*Wisteria floribunda*) vines are associated with the expression of conserved *KNOX* genes"

### Supplemental Figure 1

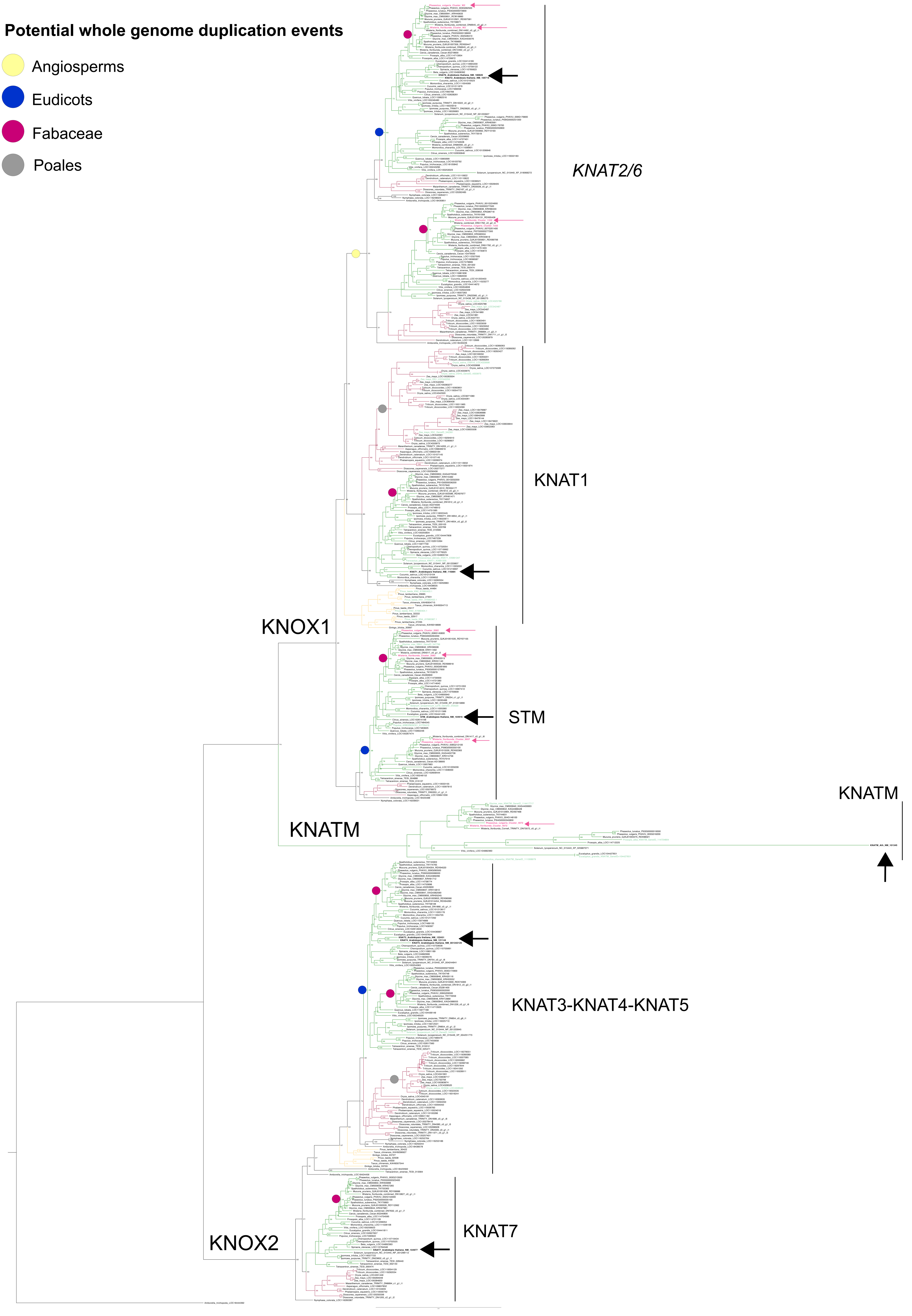

### Supplemental Figure 2

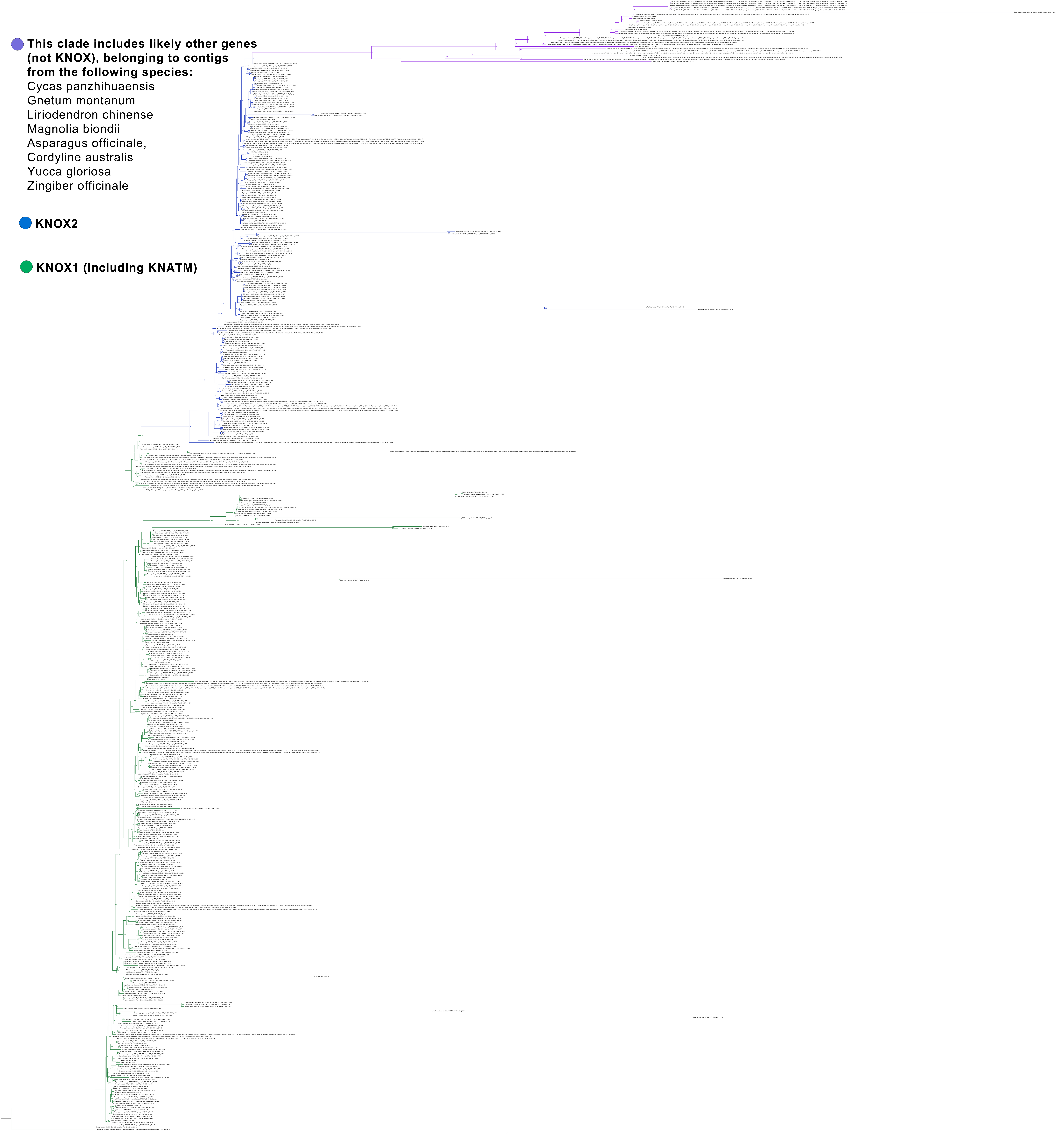

### Supplemental Figure 3a

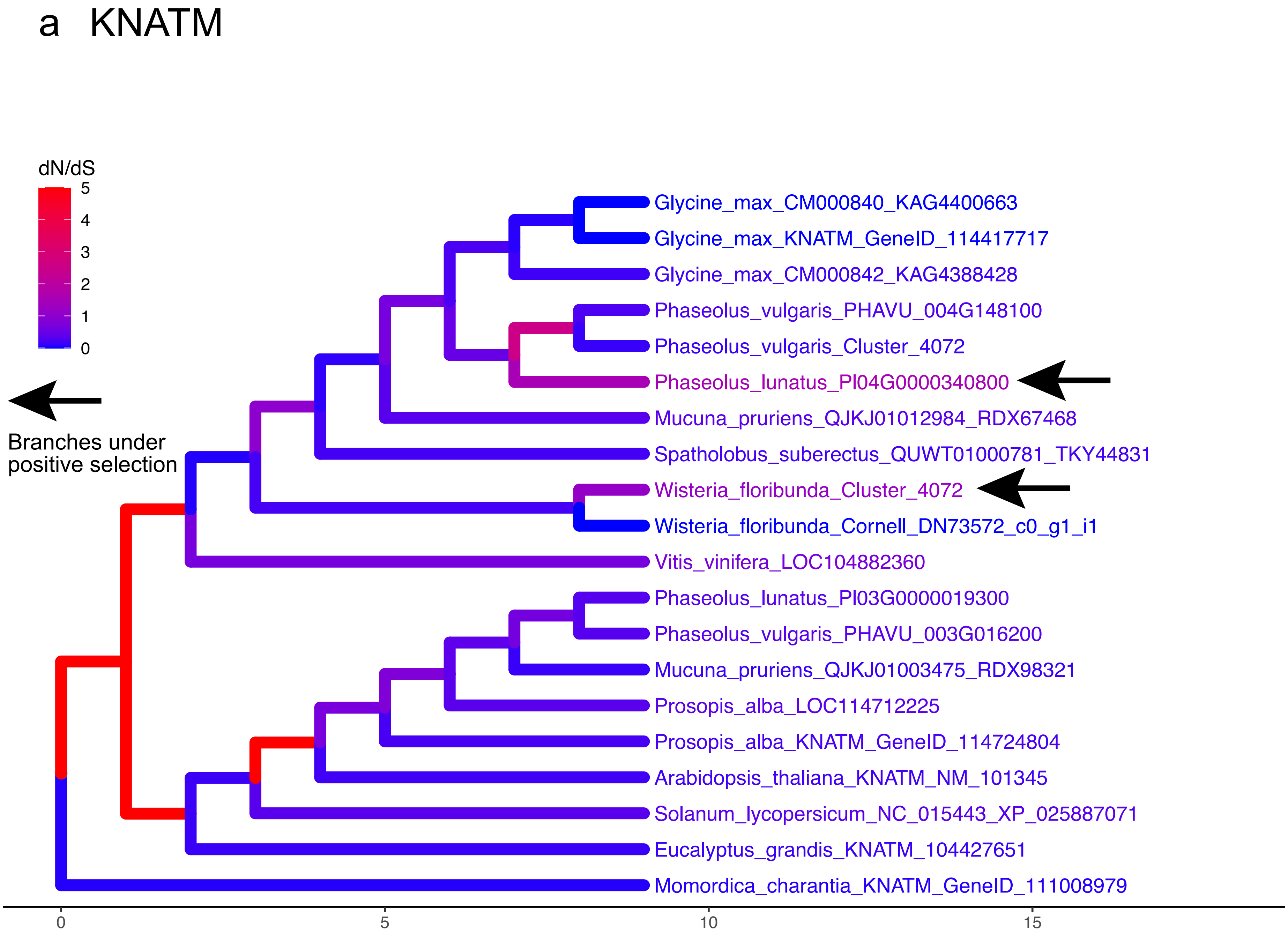

### Supplemental Figure 3b

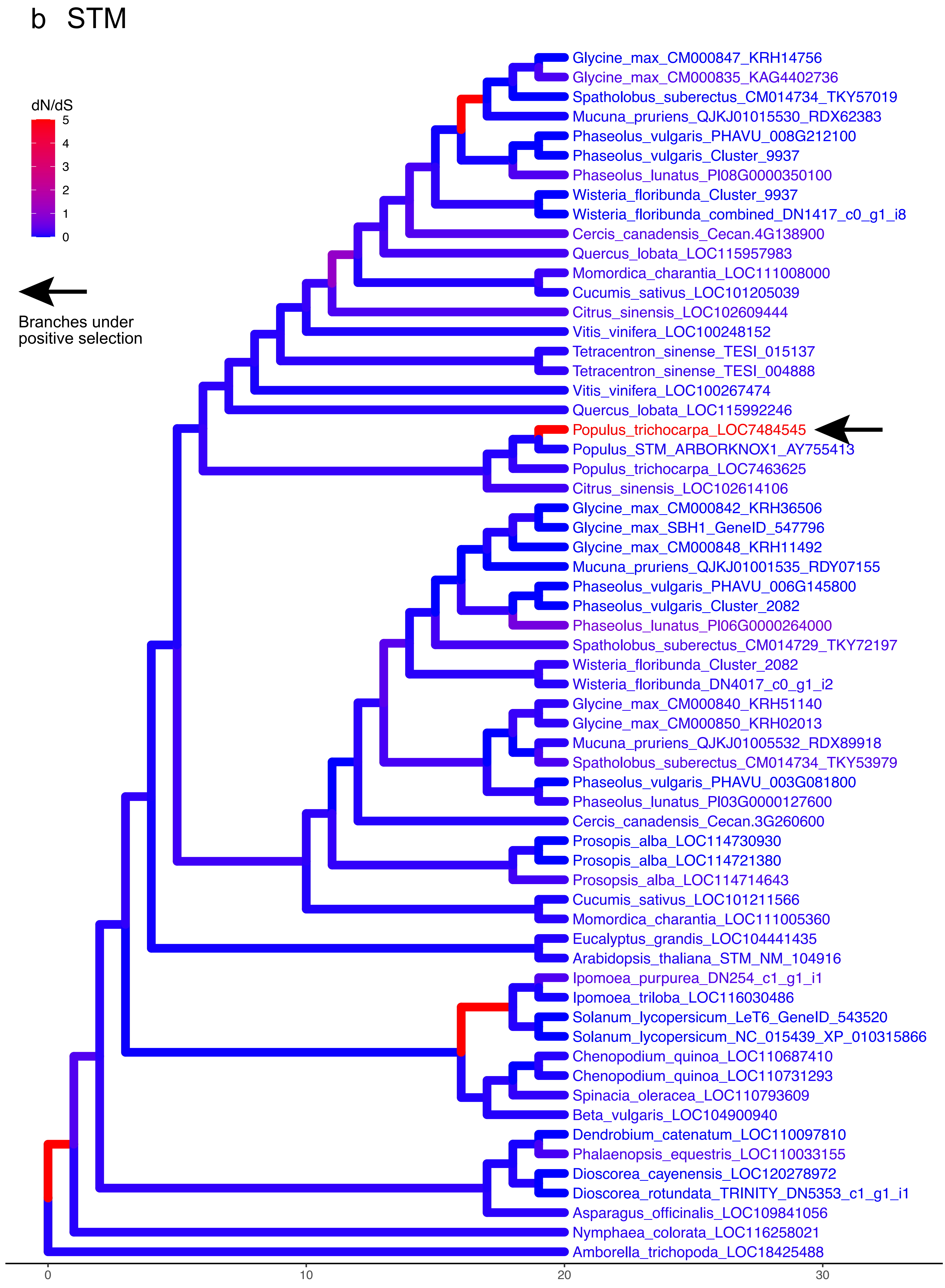
